# Electron microscopic observation of infected CCB and whole genome analysis of Koi herpesvirus isolate GY01

**DOI:** 10.1101/2020.05.21.107482

**Authors:** Hongan Duan, Ye Xu, Yi Zhou, Fengzhi Wang, Chao Ding, Jie Cao

## Abstract

**Background:** *Cyprinid herpesvirus-* 3 (CyHV-3), commonly known as Koi herpesvirus (KHV) can induce infectious and acute viremia in common/koi carp (*Cyprinus carpio*). In an earlier study in this laboratory a KHV isolate GY1506 (KHV GY01 in GeneBank) was isolated from diseased common carp, replicated on CCB cells and identified by PCR targeted on and phylogenetic analysis of thymidine kinase (TK) gene. Electron microscopy examination of GY01 infected CCB cell line and whole genome analysis was studied for further characteristics and epidemiological features of this strain.

**Results:** Electron microscopy examination of CCB infected with KHV GY01 strain revealed destruct of infected cells including incomplete nuclear membranes, deformed nucleus, marginalized nuclear chromosome, and the virus of different development stages and morphologies in the cytoplasm and nucleus, which resembles the herpesvirus. MEGA X and phyML were used for multiple alignment and phylogenetic analysis of whole genomes of GY01 and other 21 KHV strains available in GeneBank. analysis showed that GY01 was more close to E and KHV-I and was predicted it originated from the same ancestor as the E and KHV-I. Pairwise alignment of strain GY01 and strain E by Geneious software and YASS online version revealed that two strains had high identity(99.1%) at the nucleotide sequence level although variations and disagreement existed. The number and structure arrangement of open reading frames (orfs) or protein-encoding genes of GY01 is very similar to KHV E, and also to KHV-U but different from KHV-I. The characteristics and function of each orf need further study in the future.

**Conclusions:** Pathogenic changes of infected CCB cells and morphologies of KHV GY01 resembles the herpesvirus. Pairwise, multiple alignment and phylogenetic analysis of whole genomes of GY01 and other 21 KHV strains available in GeneBank demonstrated that the GY01 is closely related to strain KHV E and KHV-I and suggested it originated from the same ancestor as the E and KHV-I.

## Introduction

*Cyprinid herpesvirus-* 3 (CyHV-3), commonly known as Koi herpesvirus (KHV) is an aetiological agent for koi herpesvirus disease or infection of Koi herpesvirus that can induce infectious and acute viremia in common/koi carp (*Cyprinus carpio*).CyHV-3 and other cyprinid herpesviruses: CyHV-1 (carp pox virus, fish papilloma virus) and CyHV-2 (goldfish haematopoietic necrosis) virus were classified into the members of the genus Cyprinivirus, the family Alloherpesviridae(Davison et al., 2013;OIE, 2018). Koi herpesvirus disease appeared in the late 1990s, and has spread rapidly around the world. Gill lesions, patches and ulcerations of skin, superficial haemorrhaging at the base of the fins and /or anorexia, enophthalmia (sunken eyes) were the most reported clinical signs. But these are not pathognomic gross lesions. Confirmative diagnosis usually depend on virus isolation and identification and/or direct detection of viral DNA. Transmission electron microscopy examination of affected tissue samples and cell lines infected with diseased fish organ extracts had been tried to reveal the virus(Oh et al., 2001;Tu et al., 2004;Dishon et al., 2005;Pokorova et al., 2005). Several fish cell lines including CHSE-214, EPC, FHM and^1^ the koi fin (KF-1) were inoculated diseased fish tissue homogenates for virus isolation(Oh et al., 2001). CPE in KF-1 and transient changes in EPC was observed. Virions were detected in both gill and liver tissues. Hexagonal nucleocapsids with a diameter of 110 nm were present in nuclei or near the nuclear membrane.The capsids of negative stained virions purified from infected KF-1 cells had a diameter of about 108 nm and the virion ranged from 180 to 230 nm in diameter(Hedrick et al., 2000). Other study found the viral nucleocapsid measured at 100-110nm in diameter and surrounded by the envelope (Ilouze et al.,2011) genomes of earlier KHV type isolates from different geographic areas have been sequenced and full length genome sequences of 22 strains are available in NCBI GeneBank. The genome size of KHV strains was confirmed as 295kbp.Researches revealed that three KHV strains isolated respectively from Japan(AP008984.1, strain TUMST1(J), Israel (DQ177346.1 KHV-I), and the United States of America (USA)(DQ657948.1, KHV-U) were closely related and shared high similarity at the sequence level(more than 99%). KHV-U and KHV-I had more closer relationship than either of them to KHV J. It was postulated that they originated from same ancestor and two lineages termed as the European (U/I) and Asian(J) existed. The European lineage branched into strain KHV-U and KHV-I while strain TUMST1(J) represented the Asian lineage (Aoki et al., 2007).

A third intermediate between U/I and J was proposed by PCR targeting on two molecular markers presenting genetic variations in a study in France(Bigarré et al., 2009). Sequence analysis of two variable regions(Marker I and Marker II) of KHV isolates in Indonesia suggested a new intermediate lineage might emerge in that country (Sunarto et al., 2011).

In a later research Targeted genomic enrichment was used to recover full genomes from 1 Poland and 6 Indonesian diseased carp gill samples.Sequence analysis revealed the existence of mixed genotype infections although very low genetic diversity among specimens at the genome level(more than 99.95% of sequence identity)(Hammoumi et al., 2016)

Amplification of eight regions with the variable number of tandem repeats (VNTR) was used as a tool for KHV classification (Avarre et al., 2011) and a new TaqMan qPCR based on VNTR 3 sequence was established in 2017 to distinguish variants of Asian and European lineages(Klafack et al., 2017).

Later multiple genome alignment of 11 KHV strains were made by MAFFT online and phylogenetic tree was constructed by UPGMA(unweighted pair group method with arithmetic means) which found that strain T, J,M3 were Asian lineage while other 7 including KHV-U,E,I,KHV-I,FL,Cavoy, GZ11 and GZ11-SC were European lineages(Gao et al., 2018).

In a recent study in this laboratory a KHV strain GY1506 (available in GeneBank as KHV GY01, accession number MK260013,hereafter GY01) was isolated from diseased common carp in a farm in Jiangsu Province(WANG et al, 2017).The supernatant of the diseased fish tissue homogenate was inoculated into CCB cells. PCR was performed according to the OIE diagnostic manual, Sequence analysis of the amplified PCR product revealed its homology to thymidine kinase (TK) gene of the KHV-J strain was 100%. A phylogenetic tree constructed based on the TK gene sequence demonstrated that the strain was an Asian genotype. In this paper electron microscopy examination of GY01 infected CCB cell line and whole genome analysis in order to further investigate characteristics and epidemiological features of this isolate.

## Materials and methods

### Cell line and virus isolates

Common carp brain cell lines (CCB) was cultured with MEM (Gibico) containing 10% fetal bovine serum(FBS) at 22C.When cells grown into 80% confluent monolayer medium was emptied and inoculated with KHV GY01. After CPE appeared, the cells were harvested for electron microscopy examination and for nucleic acid purification for sequencing.

### Electron microscope examination of infected CCB cell line

The method of preparing virus-infected cell samples and ultrathin sections is similar to that reported (Dong et al., 2010).The 4^th^ passage of KHV GY01 strain were inoculated onto CCB cell lines grown in 3 flasks (25m^2^, Corning 430168).Medium was poured out and cells were scraped off after 60% CPE appeared on 12 −18 days post-inoculation (dpi), collected into centrifuge tube and centrifuged to make rice grains large, then pre-fixed with a 0.1M phosphate buffer containing 2.5% glutaraldehyde, and fixed after a 0.1 M phosphate buffer containing 2.0% osmium tetroxide. Ultra-thin sections were stained with uranium acetate-lead citrate and examined with a TEM (H-7650, 1600CCD) Hitachi transmission electron microscope (Shanghai Chenmai New Materials Testing Center).

### Sequenceing and multiple genome alignment and analysis of GY01 and other KHV strains available in GeneBank

The 6th passage (p6) of KHV GY01 strain was inoculated onto CCB cell lines. When complete CPE appeared, about 120mL cell suspensions were centrifuged at 7 000 g for 10min, ultra-centrifuged at 120,000 g for 1h, pellet was resuspended in 1mL PBS and sent to Shanghai Hanyu Biotechnology Co., Ltd. for sequencing. The genome sequence was then submitted to GeneBank. The accession number were MK260013.1,named as CyHV-3 strain GY-01.Multiple alignment and the phylogenetic tree analysis was performed on the entire genomes of all CyHV isolates available in GeneBank (table1) using MEGA X software(Kumar et al., 2018) and PhyML online execution by default options available on website http://www.atgc-montpellier.fr/phyml/.The phylogenetic relationship was inferred using the unweighted pair group method with arithmetic mean method (UPGMA),neighbor-joining(NJ), maximum likelihood(ML), maximum evolution(ME),maximum parsimony(MP) by MEGA X. The bootstrap consensus tree inferred from 1000 replicates (for ML and MP, 100replicates) was taken to represent the evolutionary history of the strains analyzed. Branches corresponding to partitions reproduced in less than 50% bootstrap replicates were collapsed. The percentage of replicate trees in which the associated strains clustered together in the bootstrap test (1000 or 100 replicates) were shown next to the branches. The evolutionary distances were computed using the number of differences method and were in the units of the number of base differences per sequence. Codon positions included were 1st+2nd+3rd+Noncoding. All positions containing gaps and missing data were eliminated.

**Table 1.** Whole genomes of KHV strains available in GeneBank

| KHV strains | Length(bp) | Submitted date | country/area isolated |
| --- | --- | --- | --- |
| KP343683.1CyHV 3 FL BAC revertant ORF136 Luc | 297,726 | 2-Jan-15 | Belgium |
| KP343684.1CyHV 3 FL BAC revertant ORF56-57 Del ORF136 Luc | 294978 | 2-Jan-15 | Belgium |
| MG925487.1 CyHV 3 strain FL | 294,636 | 4-Feb-18 | Belgium |
| MG925490.1 CyHV 3 strain M3 | 295,316 | 4-Feb-18 | Belgium |
| KJ627438.1 CyHV 3 strain KHV-GZ11 | 295,119 | 25-Mar-14 | China |
| MG925488.1 CyHV 3 strain GZ11-SC | 295,053 | 4-Feb-18 | China |
| MG925491.1 CyHV 3 strain T | 295,104 | 5-Feb-18 | China |
| MK260013.1 CyHV 3 strain GY-01 | 295,736 | 3-Dec-18 | China |
| KX544843.1 CyHV 3 isolate J2_101110 | 294,652 | 12-Jul-16 | Indonesia |
| KX544844.1 CyHV 3 isolate CB4_181110 | 294,604 | 12-Jul-16 | Indonesia |
| KX544845.1 CyHV 3 isolate J1_101110 | 294,611 | 12-Jul-16 | Indonesia |
| KX544846.1 CyHV 3 isolate I_09_2i3 | 295,092 | 12-Jul-16 | Indonesia |
| KX544847.1 CyHV 3 isolate I_10_3 | 295,142 | 12-Jul-16 | Indonesia |
| KX544848.1 CyHV 3 isolate PP3_070411 | 294,467 | 12-Jul-16 | Indonesia |
| DQ177346.1 KHV-I | 295,138 | 22-Aug-05 | Israel |
| MG925489.1 CyHV 3 strain I | 295,029 | 4-Feb-18 | Israel |
| MG925485.1 CyHV 3 strain Cavoy | 294,645 | 4-Feb-18 | Israel |
| AP008984.1 CyHV 3 strain TUMST1(J) | 295,052 | 26-May-05 | Japan |
| KX544842.1 CyHV 3 isolate PoB3 | 295,085 | 12-Jul-16 | Poland |
| MG925486.1 CyHV 3 strain E | 294,978 | 4-Feb-18 | UK |
| DQ657948.1 CyHV 3 strain KHV-U | 295,146 | 23-May-06 | USA |
| NC_009127.1 CyHV 3 (identical to KHV-U) | 295,146 | 15-Mar-07 | USA |

### Pairwise alignment of strain GY01 and E

Pairwise alignment of strain GY01and E was conducted by Geneious alignment(www.geneious.com)and YASS (https://bioinfo.lifl.fr/yass/yass.php).Geneious alignment options were set as cost matrix=65% similarity(5.0/-4.0),gap open penalty=12,gap extension penalty=3, alignment type=global alignment with free end gaps and automatical determination of sequences’ direction.

YASS (genomic similarity search tool) options were as follows: Scoring matrix (match, transversion, transition, other)=+5,-4,-3,-4(composition bias correction), Gap costs (opening, extension)=-16 and −4,E-value threshold=10,X-drop threshold=30.Other parameters were set as default.

### Orf layout of strain GY01 and strain E

Based on the multiple alignment and phylogenetic trees Genome sequence of strain GY01 and strain E were chosen and uploaded to Unipro Ugene(Okonechnikov et al., 2012) to display the orf structure. The orf layout were saved and compared.

## Results

### Electron microscope examination of infected CCB

Electron microscopy of CCB infected with KHV GY01 found that some nuclear membranes were incomplete, the nucleus was deformed, the nuclear chromosome was marginalized, and the immature virus arranged in a lattice shape in the cytoplasm and nucleus(figure 1). There were a large number of mature and immature virus particles. Three types of virus can be seen: hexagonal hollow structure, round capsule with internal electron dense substance and a clear visible structure of capsule, capsid and core. A virus with or without a capsule in the CCB infected with GY01 is about 90nm X110nm. A clear hexagonal virus is visible at the edge of the nuclear membrane at the arrow. The cytoplasm protrudes into a bubble (figure 2B).

**Figure 1.**
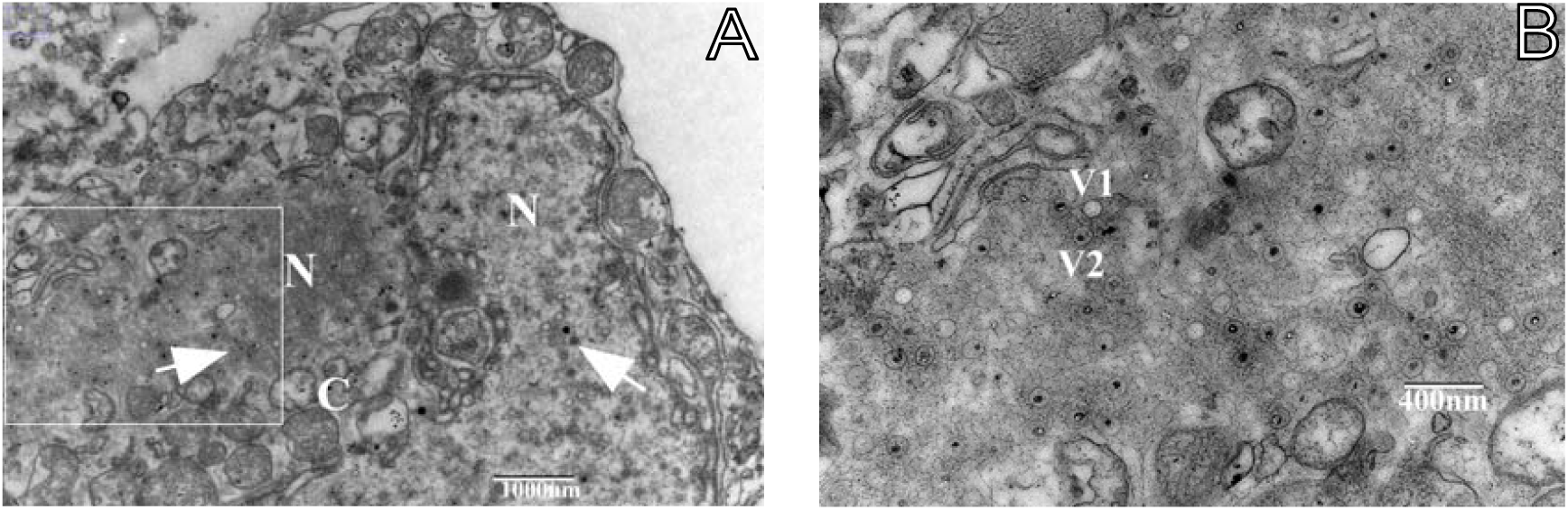
TEM micrographs of CCB infected with KHVisolate GY01 on 18 dpi. A(left): enlarged nucleus of infected CCB containing large number of various virus capsids and marginalized cytoplasm organelles. B (right) higher magnification of the left-side square of picture A showing electron lucent (V1) and electron dense(V2) capsid-like structures. *N* = Nucleus; *C* = Cytoplasm; Arrow = various kind of capsids.

**Figure 2.**
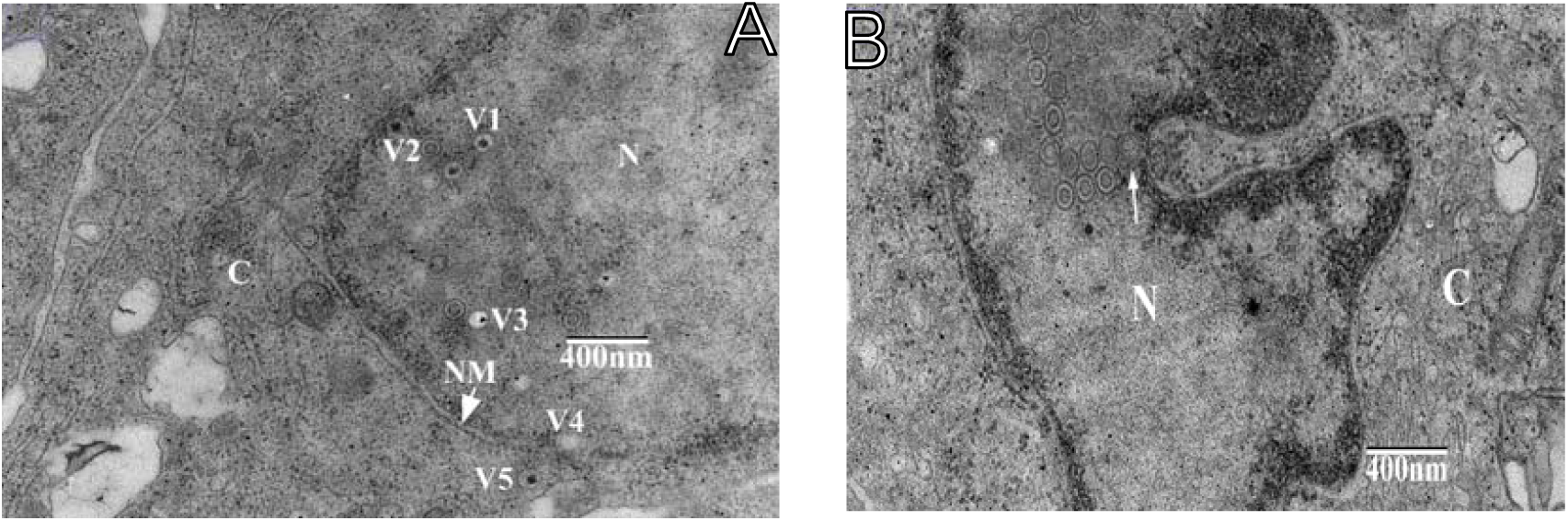
TEM micrographs of CCB infected with KHV isolate GY01 on 18 dpi. (A) various kind of capsids inside and outside of infected CCB nucleus. (B) a clear hexagonal viral capsid (arrow)is visible at the edge of the nuclear membrane. The cytoplasm protrudes into nucleus forming a bubble. *N* = Nucleus; *C* = Cytoplasm; NM=nucleus membrane; V1,V3= electron dense(below the letter V1) and electron lucent(left of the letter V3) capsid-like structures (capsids);V2= Concentric circle-like (right of the letter V2)capsids.V4, V5=virus particles near and outside the nucleus membrane, Arrow in B = various kind of capsids.

### whole genome sequencing and analysis

Genome common characteristics of Strain GY01 are listed in table2 and table3

**Table 2.** Strain GY01 Genome common characteristics

|  |  |
| --- | --- |
| Length: | 295 736 |
| GC Content: | 59.24% |
| Molar Weight: | 91414358.08 Da |
| Molar Ext. Coef: | 3118100500 l/mol |
| Melting TM: | 89.18□ |
| nmole/OD260 : | Non-applicable |
| μg/OD260 : | 29.32 |

**Table 3.**
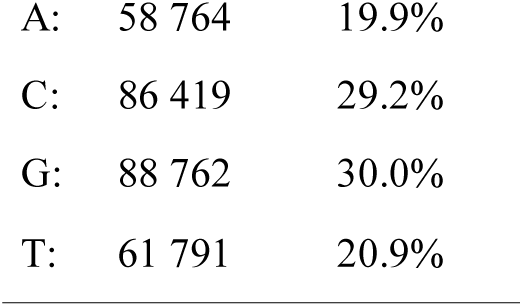
Characters occurrence

GY01 formed a separate branch when the evolutionary history was inferred by using UPGMA (figure 3A), NJ and MP method(not shown) by MEGA X. KHV-U is identical to NC009127 and more close to KHV I than to KHV-I in UPGMA tree. while GY01 is more closely related to strain J in ME and ML tree. PhyML tree gave similar results to the UPGMA tree (figure 3B).

**Figure 3.**
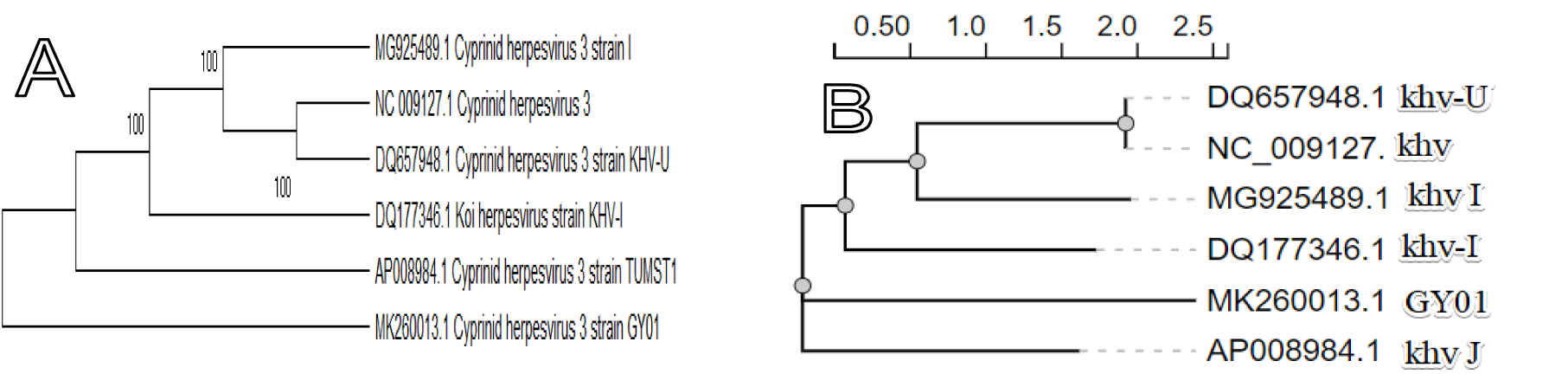
Evolutionary relationships of GY01 and KHV-U, KHV-I and J strains. The evolutionary history was inferred using the UPGMA method (A) and with the bootstrap consensus tree inferred from 1000 replicates and branches corresponding to partitions reproduced in more than 50% bootstrap replicates and PhyML (B).

12 KHV strains were classified into 4 groups. KHV T, GZ11 and KHV I and KHV-U belonged to lineage 1a, 1b included GY01, KHV E and KHV-I, lineage 1c consisted KHV M3 and KHV J. lineage 2 included KHV FL and Cavoy isolated from Belgium and Israel respectively. GY01 was close to E and KHV-I while M3 and J, FL and Cavoy, KHV-U(NC_009127) and KHV I(figure 4A) were most close pairs in UPGMA tree, similar results acquired from ME,MP and ML. GY01 looked like forming a separate branch in NJ tree actually it was still closer to KHV-I and E(data not shown). PhyML tree categorized 12 KHV strains into 3 lineages(groups), KHV GZ11 and KHV-U(NC009127) each formed a branch respectively while other 10 formed a big group in which GY01 was more closely related to KHV E then KHV-I (figure 4B).

**Figure 4.**
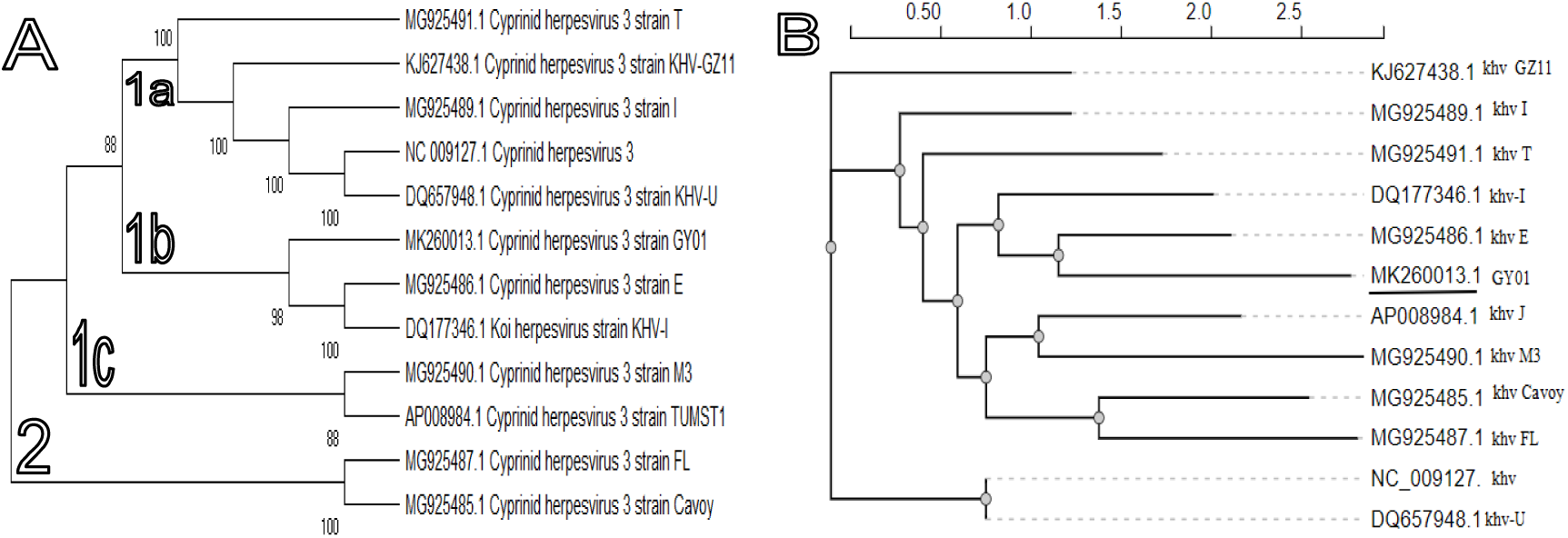
Evolutionary relationships of GY01 and other 11 strains. The evolutionary history was inferred using the UPGMA method (A) and with the bootstrap consensus tree inferred from 1000 replicates and branches corresponding to partitions reproduced in more than 50% bootstrap replicates and PhyML (B).

In figure 5A 1a consisted strain T,I and KHV-U and GZ11 strains. While GY01 belonged to 1b which had KHV-I, E and 2 Indonesia strains. Strain J and M3 in 1c were closely related to each other.1c included Cavoy and PoB3 which was isolated in Israel and Poland respectively. 2a including 4 isolates from Indonesia and 3 FL derivatives. In figure 5B lineage 1a consisted 6 isolates from Indonesia.1b was a big group. lineage 2 and 3 was KHV M3 and KHV J respectively. GY01 was closely related with KHV-I and E.

**Figure 5.**
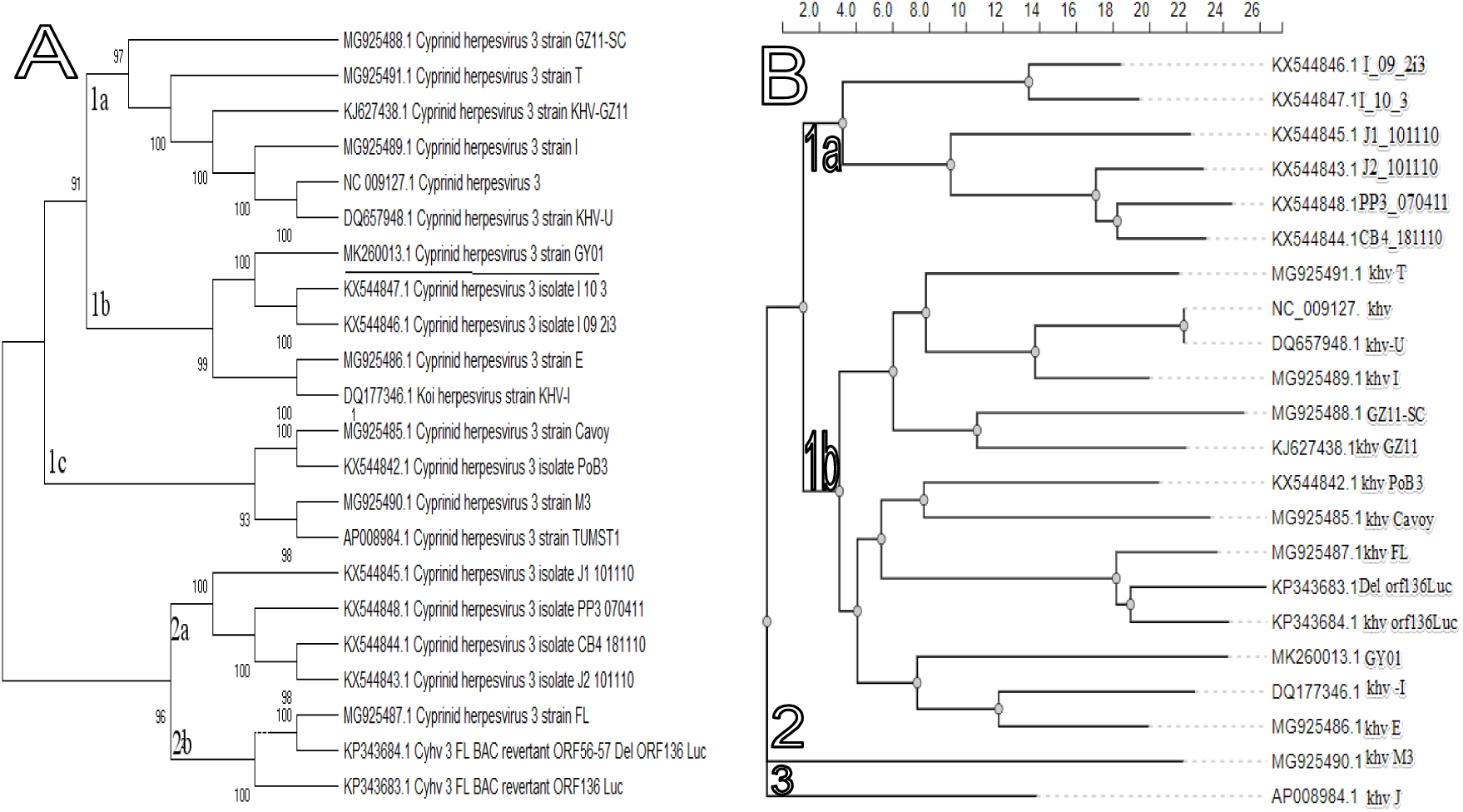
Evolutionary relationships of GY01 and other 21 strains. The evolutionary history was inferred using the UPGMA method (A) and with the bootstrap consensus tree inferred from 1000 replicates and branches corresponding to partitions reproduced in more than 50% bootstrap replicates and PhyML (B)

**Figure 6.**
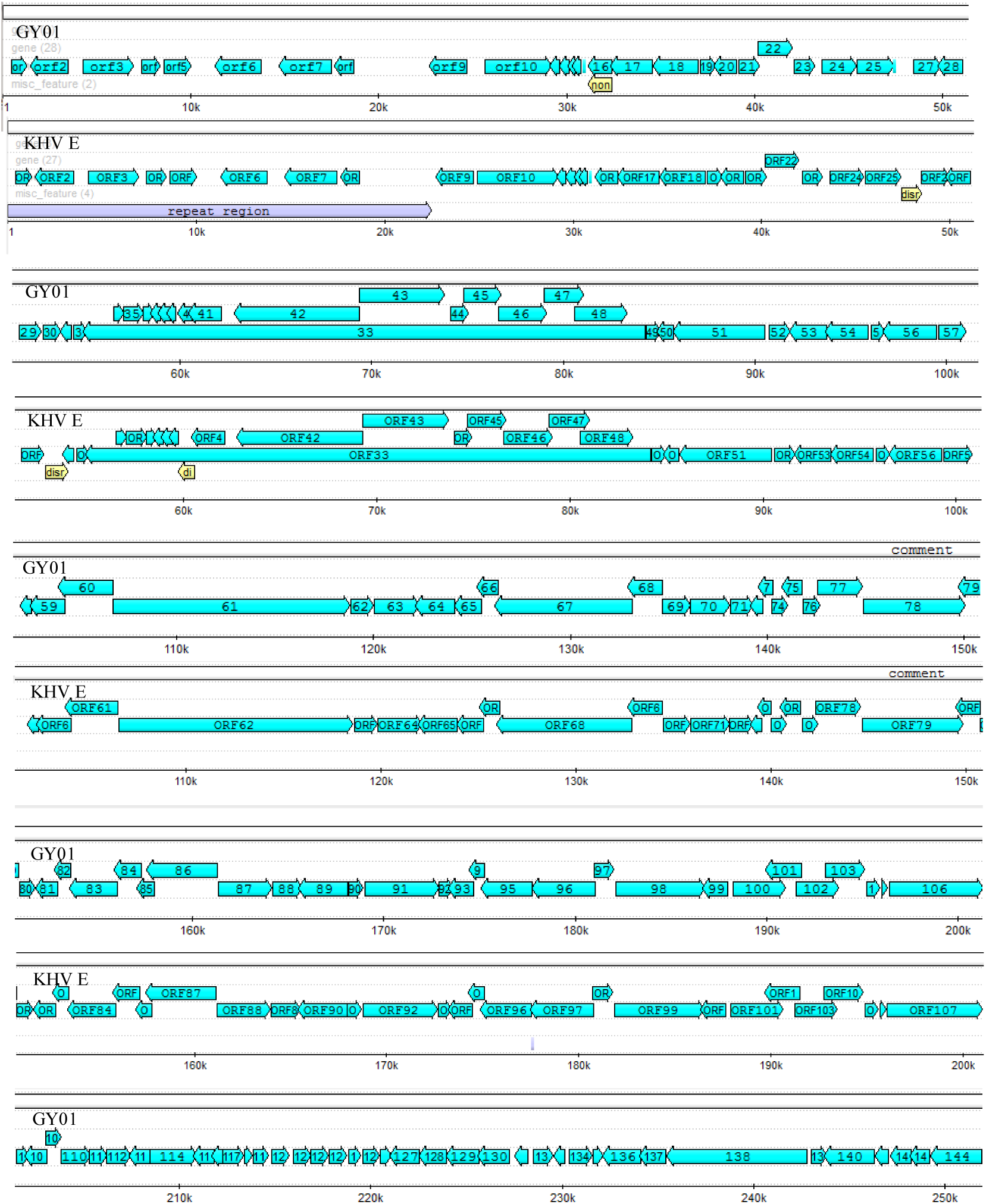
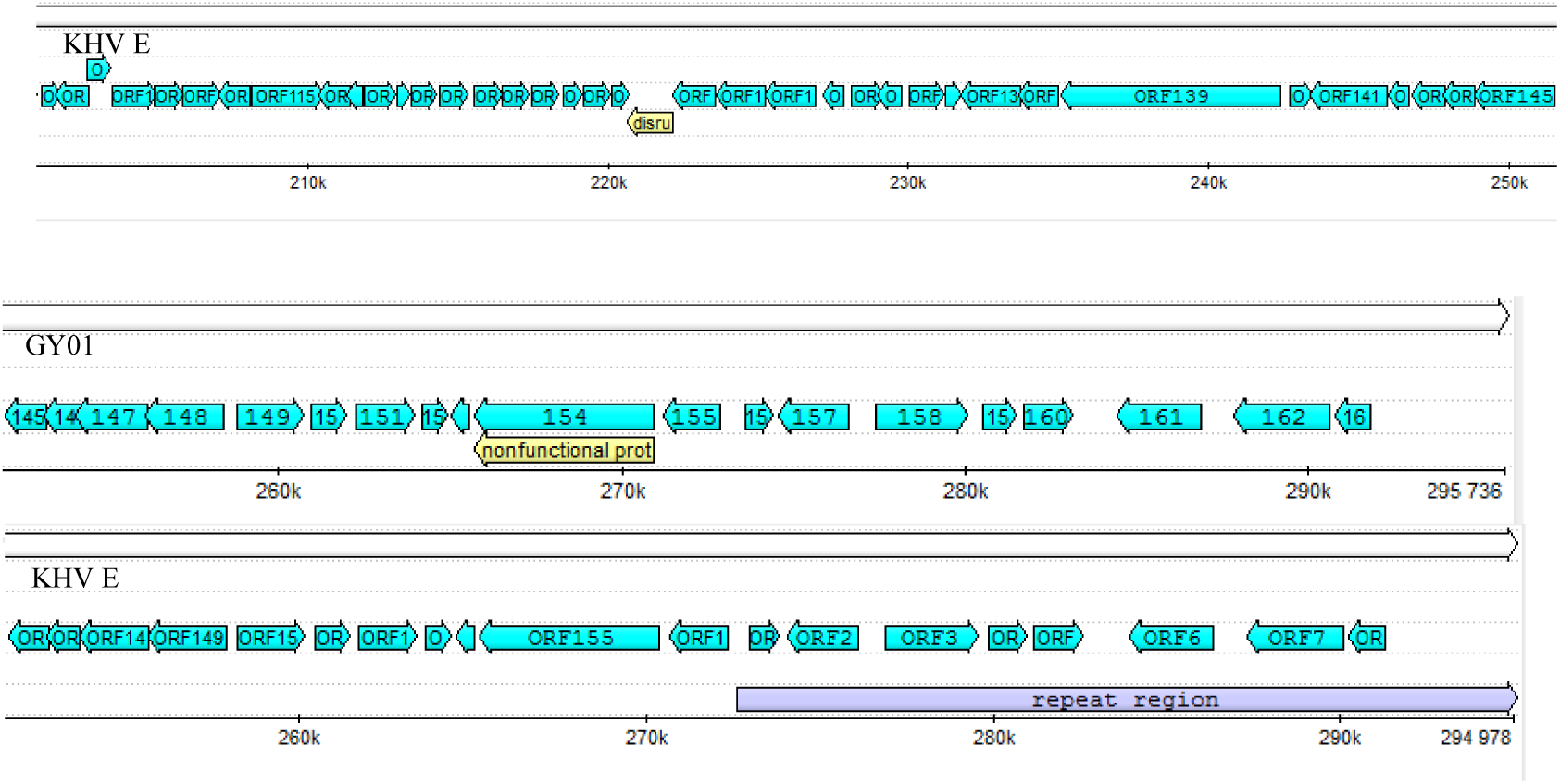
Open reading frame (orf) layout of strain GY01 and KHV E. orf: open reading frame (orf). non: non-functional protein coding region. Dis: disrupted orf.

### Pairwise alignment of strain GY01 and strain E

Geneious analysis showed that identical sites of strain GY01 and KHV E were 293,820 (99.1%) with 99.1% Pairwise Identity. Mean number of ungapped lengths of 2 sequences was 295357.0 with Std Dev= 379.0, Minimum= 294978 and Maximum= 295736.

There were 574 disagreements including ambiguous, indels and gaps, of which 341 were single nucleotide variations and 80 were two or three base differences.

There were 3104 alignments found by the YASS, and three long fragments were with high identities (more than 98%) are listed in table 4 and gaps less than 0.01%. While other shorter fragments were not listed here. 22kb long terminal repeats of KHV GY01 were present at both ends of genomes as that of KHV E.

**Table 4.**
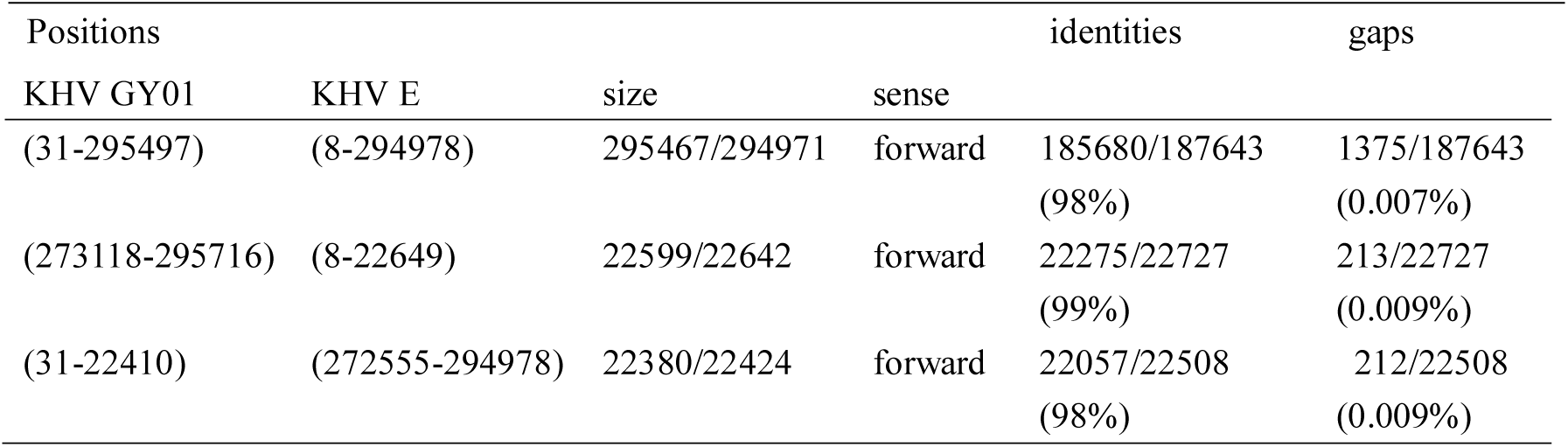
Long fragments of strain GY01 and E with high similarity

GY01 genome had 163 open reading frames (orfs) of which orf 16 and orf 154 was annotated as nonfunctional protein coding regions. KHV E genome had 156 orfs numbered orf 1 to orf 156 with duplicate orf 1 to orf 8 at the both end of genome while orf 26, orf 30, orf 40, orf 58 and orf 128 disrupted. At the both end of genome existed a long repeat region, i.e. orf1 to orf 8 duplicate which was similar to the orf 156 to orf 163 at the right end of GY01 genome.

## Discussion

### Morphology of cell line infected with KHV GY01 by electron microscopy examination

In this study Electron microscopy examination of CCB infected with KHV GY01 strain found that pathogenic changes of infected cell structure including incomplete nuclear membranes, deformed nucleus, marginalized nuclear chromosome, and the virus of different development stages and morphologies in the cytoplasm and nucleus (figure 1) were similar to previous researches. Three types of virus particles could be seen: Concentric circle, electron dense and electron lucent-like capsids or virus particles. The cytoplasm protruded into a bubble (figure 2B).This phenomenon and existence of capsids in and out of nucleus membrane suggests the replication of viral genome in nucleus and viral particle development further in the cytoplasm. Anyhow complete viral particle with envelope was not found in this study probably due to the different stages of virus formation. A virus with or without a capsule in the CCB infected with KHV is about 90nm x 110nm. An earlier study in Korea found that virus-like particles of 70–80 nm in diameter were detected in FHM (fathead minnow) cells inoculated with kidney and spleen filtrate of moribund fish infected by koi herpesvirus (Oh et al., 2001) but other fish cell lines: CHSE-214(chinook salmon embryo), EPC (epithelioma papulosum cyprini), and RTG-2(rainbow trout gonad) had no CPE or transient CPE on EPC. Electron microscopy observation of KHV virion formation on three cell lines of common carp revealed similar cell destruction and capsids of about 110 nm in diameter with three different types of nucleo-capsid which were a capsid containing an internal ring (spherical), structure containing electron dense material and empty shell. Mature enveloped virions of 170-200 nm in diameter existed in cytoplasm or outside the cells (Miwa et al., 2007). In an another report the morphology of KHV virus was simply divided into enveloped and non-enveloped virion(Dong et al., 2011). Ultrastructural analysis of gill tissues sampled from KHV infected Nile tilapia found that viral particles **w**ith a diameter of 150-170 nm in the cytoplasm or outside the gill cells(Wahidi et al., 2019).

Different size of capsids or virion particles in diameter in above researches probably due to the different cutting side of section or different stages of virus development but the morphologies were similar and consistent with viruses from the Herpesviridae family.

### Multiple alignment and phylogenetic analysis of GY01 and other KHV strains available in GeneBank

In this study two software i.e. MEGA X and phyML were used for multiple alignment and phylogenetic tree construction for GY01 and other genome sequences of KHV strains available in GeneBank. For most strains MEGA X and phyML gave similar results even though differences among methods contained in MEGA X and phyML existed for some strains.

NC_009127 is identical to KHV-U and served as internal control. KHV-U(NC_009127) is closer to KHV I other than KHV-I. While in most references KHV-U and KHV-I are referred as European genotype (U/I). M3 and J, Cavoy and FL/PoB3 are most closely related strains to each other.KHV T here was in a lineage including KHV-U,I and GZ11 which was regarded as European genotypes although strain T was often classified as Asian lineage which including M3 and J explained later in this article. As for the six Indonesian strains UPGMA tree showed that 4 of which were close to strain FL and their derivatives and other two (I_10_3 and I_09_2i3 closer to KHV E, KHV-I and GY01, while phyML tree revealed that the six Indonesian strains formed 1 separate sub-lineage.

Origin and genotypes or lineages of KHV strains were studied and predicted in two directions: whole genome analysis and specific short regions of KHV genomes such as variable markers or a certain gene for an example TK gene.Two lineages were proposed as Asian and European after whole genome of three KHV strains isolated from Japan (KHV J), Israel (KHV-I), and USA(KHV-U) were compared (Aoki et al., 2007). The existence of two lineages was confirmed by a duplex PCR targeting at two molecular markers with genetic variations and PCR product sequencing from 42 samples of infected carps from France, The Netherlands and Poland. This method also identified a third genotype or genetic intermediate between U/I and J (Bigarré et al., 2009).

Three enlarged known PCR target regions of Sphl-5, 9/5 and the thymidine kinase (TK) gene of KHV genome were amplified by PCR from 43 samples including the reference strain of the Japanese KHV from Asian countries/areas except Israel and 16 from outside Asian countries including one from the USA and one from Israel. Based on comparison of the PCR product sequences of ten variable areas of genome, KHV strains were classified into Asian and European genotypes, which consisted of 2 (A1-A2) and 7 variants (E1-E7) respectively. While European genotypes including strains from USA and Israel (Kurita et al., 2009). The reference KHV strains from Japan, USA ang Israel in above study were probably the strains KHV J, KHV-U and KHV-I although the origin or GeneBank Accession Number could not be traced back.

Phylogenetic analysis by the Neighbor-Joining methods using the MEGA 4.1was conducted (Sunarto et al., 2011) on TK gene sequences of 4 KHV isolates from USA, UK and Indonesia and 14 ethanol-fixed tissue samples in Indonesia as well as sequences of KHV -U (USA), J (Japan) and KHV-I (Israel), European and Asian genotypes available from GenBank. All TK gene sequences were branched into European and Asian genotypes which were consistent with the previous report (Kurita et al. 2009;Aoki et al., 2007). TK gene sequences of the two Indonesian isolates and samples were identical to that of the Asian genotype. As reported previously, KHV-U was European genotype while KHV J (AP008984) and KHV-I(DQ177346) were more close to A1 than to A2 and the European genotypes. Classification of KHV-I in above mentioned study contradicted previous reports. Alignment of two variable regions (marker I and II) revealed a new allele (I++II-) in one UK and all Indonesian isolates which were not reported previously among Asian or European genotypes may represent a genetic intermediate between the two lineages. Existence of a mixture of alleles in some Indonesian isolates predicted that an individual sample may contain two different variants of KHV.

CyHV-3 Genomes of 7 specimens from Poland and Indonesia as listed in table1 in present study were recovered by targeted genomic enrichment and were aligned with three type KHV strains KHV-U,KHV-I and KHV-J using Mafft program online version(Hammoumi et al., 2016). Phylogenetic tree constructed by the neighbor-Joining method by MEGA6 showed PoB3 was European lineage while all 6 Indonesian specimens were classified into Asian one and the 3 reference KHV strains had the same results as reported earlier(Aoki et al., 2007).

Improved Phylogenetic method based on variable number of tandem repeats (VNTR) and a VNTR-3 qPCR and whole genome sequences was applied for virus typing including type KHV strains KHV-U(NC_009127),KHV-I and KHV J, KHV T, GZ11, FL-BAC (Klafack S.et al 2017). VNTR results were comparable and confirmed the results of VNTR-3 qPCR and whole genome comparison. KHV J and T formed one cluster, KHV-U, KHV-I and FL-BAC were in the same group while GZ11 are inconsistent in three trees and was interpreted as an intermediate status between European and Asian lineages.

The phylogenetic tree based on fullClength genome sequences of 11 KHV strains was built using UPGMA in MEGA6 found that T, J and M3 are in Asian lineage, while strain KHV-U, E, I, Cavoy, KHV-I, FL, GZ11 and GZ11-SC in European Lineage(Gao et al., 2018). Our earlier research on comparison of TK gene of KHV strains showed that the strain GY01 close to the Asian lineage while in this report the GY01 was closer to KHV E and KHV-I which were normally regarded as European lineage. GY01 was predicted it originated from the same ancestor as the E and KHV-I.So evolutionary prediction based on specific short regions such as certain variable regions or a specific genes as well as small number of genomes may give different results compared to multiple comparison and phylogenetic analysis of a large number of whole genomes. Prudence is needed to come to a conclusion when short regions or few whole genomes are used to evolution inference for a specific strain Comparison of methods used for the whole genome alignment demonstrated that UPGMA,NJ and ME were very fast while ML, MP by MEGA X took few days to run the program and probably got no results and warning appeared “not enough capacity to run the program”. while phyML took less time to get the results and it was believed the reliable method for phylogenetic construction of the whole genomes to be analyzed. Pairwise alignment of strain GY01 and strain E by Geneious software and YASS online version showed that two strains had high identity(99.1%) at the level of nucleotide sequence although variations and disagreement existed. the number and structure arrangement of orfs or protein-encoding genes of GY01 was very similar to KHV E, and also to KHV-U but different from KHV-I (data not shown). The characteristics and function of each orf need further study in the future.

## Conclusion

Pathogenic changes of infected CCB cells and morphologies of KHV GY01 resembles the herpesvirus. Pairwise, multiple alignment and phylogenetic analysis of whole genomes of GY01 and other 21 KHV strains available in GeneBank demonstrated that the GY01 was closely related to strain KHV E and KHV-I and suggested it originated from the same ancestor as the E and KHV-I.

## Acknowledgements

This work was financially supported by the science and technology project of Jiangsu Entry-Exit Inspection and Quarantine Bureau (No. 2017KJ01) and of Nanjing Customs of P.R. China (No.2019KJ16).

